# The BD2 domain of BRD4 is a determinant in EndoMT and vein graft neointima formation

**DOI:** 10.1101/509414

**Authors:** Mengxue Zhang, Bowen Wang, Go Urabe, Yitao Huang, Jorge Plutzky, K. Craig Kent, Lian-Wang Guo

## Abstract

**Background:** Vein-graft bypass is commonly performed to overcome atherosclerosis but is limited by high failure rates, principally due to neointimal wall thickening. Recent studies reveal that endothelial-mesenchymal transition (EndoMT) is critical for vein-graft neointima formation. BETs are a family of Bromo/ExtraTerminal domains-containing epigenetic reader proteins (BRD2, BRD3, BRD4). They bind acetylated histones through their unique tandem bromodomains (BD1, BD2), facilitating transcriptional complex formation and cell-state transitions. The role for BETs, including individual BRDs and their unique BDs, is not well understood in EndoMT and neointimal formation.

**Methods and Results:** Repression of BRD4 expression abrogated TGFβ1-induced EndoMT, with greater effects than BRD2 or BRD3 knockdown. An inhibitor selective for BD2 in all BETs, but not that for BD1, blocked EndoMT. Moreover, expression of a dominant-negative BRD4-specific BD2 fully abolished EndoMT. Concordantly, BRD4 knockdown repressed TGFβ1-stimulated increase of ZEB1 protein – a transcription factor integral in EndoMT. In vivo, lentiviral gene transfer of either BRD4 shRNA or dominant negative BRD4-specific BD2 mitigated neointimal development in rat jugular veins grafted to carotid arteries.

**Conclusions:** Our data reveal the BD2 domain of BRD4 as a determinant driving EndoMT *in vitro* and neointimal formation *in vivo*. These findings provide new insight into BET biology, while offering prospects of specific BET domain targeting as an approach to limiting neointima and extending vein graft patency.

## Introduction

Endothelial-mesenchymal transition (EndoMT) is a process involving endothelial cell (EC) switching to a mesenchymal-like cell state. EndoMT is now implicated in a growing number of disease conditions ^1^. In addition to atherogenesis ^2^, EndoMT was recently identified as crucial for the failure of vein grafts ^3^, the bypass conduit surgical treatment of atherosclerosis in coronary and peripheral arteries. Despite their extensive use, vein graft failure is common, reaching 40% within 12-18 months after grafting ^3^. Thus, a compelling rationale exists for understanding the regulation of EndoMT ^1^, which might allow for novel therapeutic options in an area with limited if any progress.

In response to specific extracellular stimuli, including transforming growth factor (TGFβ1) exposure, ECs undergo a cell state transition, losing EC markers while manifesting proteins like smooth muscle actin (αSMA), calponin, and collagen-I that characterize smooth muscle cells (SMCs) ^3^. Subsequent execution of EndoMT involves distal specific transcription factors, including ZEB1 (Zinc Finger E-Box Binding Homeobox), Snail, and Twist ^1^. This EndoMT transcriptional reprogramming manifests as acquired non-EC phenotypes, including collagen production and cell migration ^1^. EndoMT remains poorly understood, in contrast to epithelial-mesenchymal transition (EMT). While EndoMT and EMT overlap in some mesenchymal-like signaling pathways and cellular characteristics ^1^, distinct EndoMT mechanisms are increasingly being recognized ^2, 3^. Despite this progress, there has been limited insight into how context-dependent epigenetic regulations influence EndoMT programs ^1^.

The Bromodomain and ExtraTerminal domain-containing epigenetic reader proteins (BETs) have been recently identified as powerful determinants of differentiation and cell-state transitions ^4, 5^. BETs are comprised of family members BRD2, BRD3, BRD4, and BRD-t (testis-specific)^5^, each containing two bromodomains (BD1 and BD2) that bind to acetylated lysines as marked on histones or present in transcription factors. Depending on cell types and environmental cues, BETs facilitate assembly of specific transcription factors with core transcription machinery to activate (or repress) a select set of genes that orchestrate cellular responses, including cell state transitions ^6^. As we and others have shown, BET inhibition can block cell state transitions involving EC inflammation, cardiomyocyte hypertrophy and SMC proliferation ^4, 7, 8^. Nevertheless, key questions remain regarding basic BET biology in vascular cells, including the relative contributions among different BET family members as well as the specific role of BD1 *vs* BD2 in BET-regulated responses. Specifically, BET involvement in neointimal formation and vein graft failure remains unexplored. Deeper understanding of BET biology may prove essential to targeting BETs for therapeutic purposes as is underway in oncology as well as heart disease ^9, 10^.

Using selective genetic and pharmacological tools applied to *in vitro* EndoMT studies as well as an *in vivo* model of vein graft neoinitimal formation, we present data here dissecting out the specific roles of individual BET family members and their bromodomains in EndoMT and neointimal formation. Our data indicate that BRD4 plays a predominant role in EndoMT over BRD2 or BRD3, effects that depend on BD2 not BD1. Further mechanistic studies reveal that BRD4 controls TGFβ1-stimulated increases in ZEB1. *In vivo*, manipulation of BRD4 or BRD4 specific BD2 in the rat vein graft model supports an essential role for this epigenetic reader protein in neointimal formation. Taken together, these studies reveal details of BET biology that determine EC functional responses, BET involvement in EndoMT underlying cell state/identity transitions, and novel potential strategies for the unmet need of preventing vein graft failure.

## Results

### Treatment with pan-BET bromodomain inhibitor JQ1 abolishes TGFβ1-induced EndoMT in rat primary aortic ECs

To investigate possible BET family involvement in EndoMT that we observed earlier^11^, we studied rat primary aortic EC responses to TGFβ1 — a canonical inducer of EndoMT ^1, 3^, doing so in the presence or absence of the pan-BET inhibitor JQ1 ^12^. JQ1 reversibly binds both BD1 and BD2 in all BETs, thus disrupting the interaction between BRD2, 3 and 4 and acetylated lysines on histone tails and certain transcription factors ^6^. EndoMT was monitored by detecting mRNA induction of three well-established EndoMT markers in ECs, namely αSMA, calponin, and collagen-I. Cells were pre-treated with vehicle or JQ1 (1 μM) prior to TGFβ1 stimulation. As expected, TGFβ1-induced EndoMT as indicated by a 2 to 4-fold induction of αSMA, calponin, and collagen-I mRNAs; in contrast, in the presence of JQ1, this cell state transition was completely abolished (Figure 1A).

**Figure 1.**
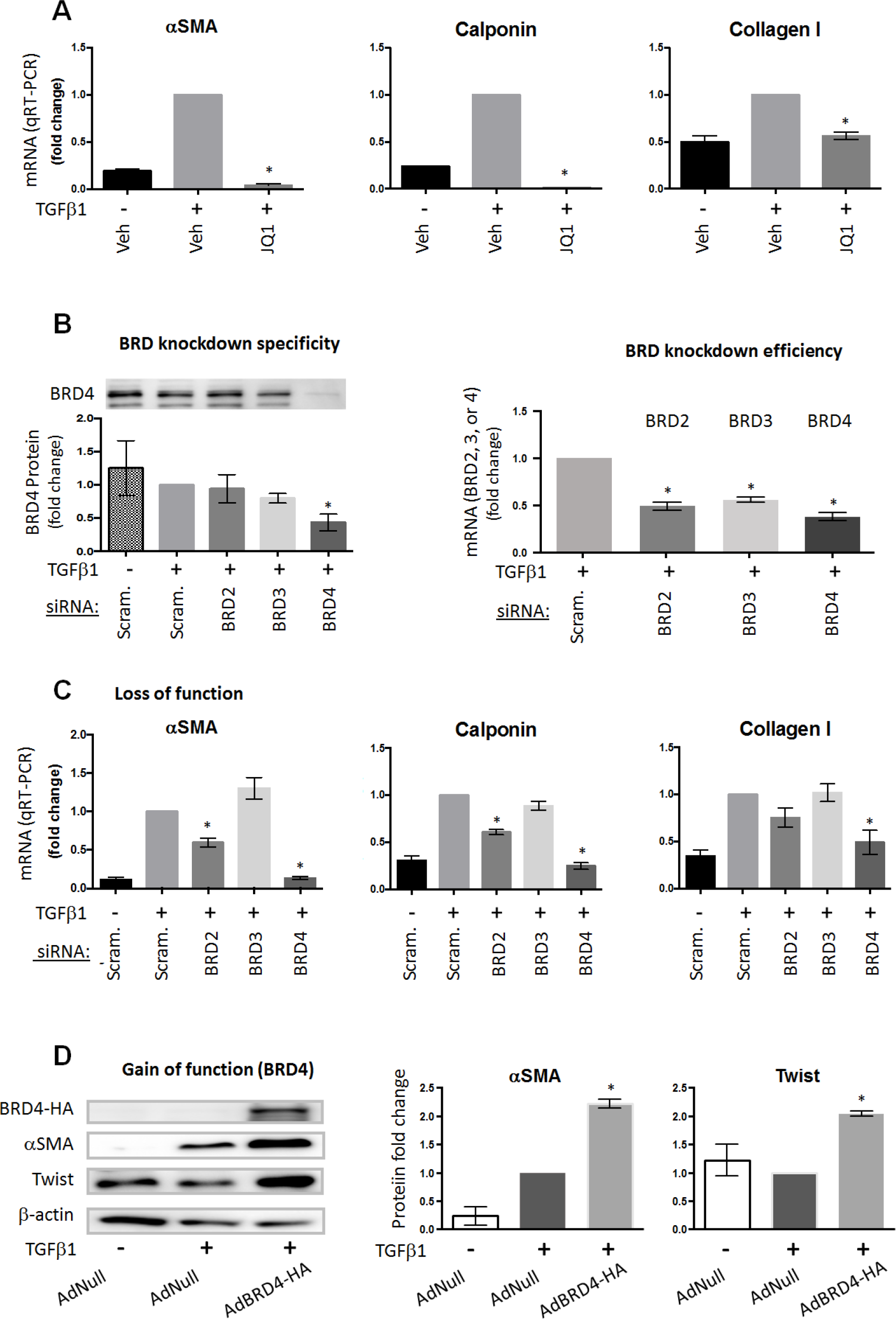
Either pan-BET inhibition or selective BRD4 silencing abrogates TGFβ1-induced EndoMT. A. The pan-BET inhibitor JQ1 represses TGFβ1-induced mRNAs of EndoMT markers. As described in detail in Methods, rat primary aortic ECs were starved for 6h in the endothelial cell culture basal medium containing 1% FBS, and then pre-treated with 1 μM JQ1 or vehicle control (equal amount of DMSO) for 2h prior to adding TGFβ1 (final 50 ng/ml). Cells were harvested at 72h after TGFβ1 stimulation and used for qRT-PCR assays. B. Efficiency and specificity of BET family member knockdown. ECs were transfected overnight with a scrambled siRNA or an siRNA specific for BRD2, BRD3, or BRD4, recovered in the complete medium for 24h, and then starved in the basal medium for 6h prior to adding TGFβ1. After 72h and 96h of TGFβ1 stimulation respectively, cells were harvested for qRT-PCR and Western blotting. C. siRNA targeting of BRD4 abolishes TGFβ1 induction of EndoMT. In the knockdown experiments (see B), the ECs stimulated (or not) with TGFβ1 for 72h were used for qRT-PCR determination of mRNA levels of EndoMT markers. D. Gain-of-function: Adeno-viral expression of BRD4-HA. ECs were transduced for 6h with AdNull (control) or AdBRD4-HA in the complete medium and recovered in fresh complete medium for 24h. Cells were then starved for 6h in the basal medium containing 1% FBS before TGFβ1 stimulation (96h) followed by Western blotting. Quantification (A-D): For each independent experiment, the control value (vehicle, scrambled siRNA, or AdNull, in the presence of TGFβ1) is presented as 1, and the values from other conditions are normalized to this control. Normalized values from independent repeat experiments (n ≥ 3) are averaged (mean ± SEM); *P<0.05 compared to the normalizing control.

### BRD4 plays a predominant role in TGFβ1-induced EndoMT

Given these potent effects of pan-BET inhibition on EndoMT, we next considered the relative contributions of BRD2, 3, or 4 in driving this EC response. As currently no BET member-specific inhibitors exist, we validated (Figure 1B) and then employed siRNAs specific to each BRD in TGFβ-induced EndoMT experiments (Figure 1C). While BRD2 knockdown reduced TGFβ1 stimulated αSMA expression by ~30%, BRD4 silencing completely blocked the action of TGFβ1, reverting EC expression of αSMA mRNA to basal levels seen in unstimulated ECs (scrambled siRNA, no TGFβ1). BRD3 knockdown showed no effect on αSMA expression, slightly but not significantly increasing its mRNA over the TGFβ1-stimulated level. Similar patterns of BRD knockdown effects on calponin and collagen-I were observed: BRD2 siRNA reduced TGFβ1-stimulated expression by ~30%, BRD3 siRNA had no effect while BRD4 siRNA effectively reduced their mRNAs to basal, unstimulated levels (Figure 1C). Further confirming the specific role of BRD4, its overexpression via adenovirus (Figure 1D) or lentivirus (Figure S1) in ECs significantly increased protein levels of αSMA, a canonical EndoMT marker, and Twist, a transcription factor involved in EndoMT ^1, 13^. Thus, while pharmacologic pan-BET inhibition blocks TGFβ1-induced EndoMT, specific genetic manipulation of individual family members indicates BRD4 as the dominant as well as sufficient BET determinant of EndoMT.

### Blocking BD2 but not BD1 of the BET family proteins abrogates TGFβ1-induced EndoMT

Since bromodomains within BETs determine their localized binding to specific acetyl modification of histone tails and subsequent transcriptional effects ^5^, we investigated whether the two highly homologous BET bromodomains ^12^ play differential roles in EndoMT. Pharmacologic BET inhibitors exist that have been reported to have selective BD-blocking effects, namely RVX208 as inhibiting BD2 and Olinone as preferentially targeting BD1 (Figure 2A), albeit without discrimination between BRD2, 3 and 4 ^14, 15^. We found that while pre-treatment with RVX208 significantly decreased TGFβ1-stimulated αSMA protein levels as compared to vehicle control (+TGFβ1), an equivalent concentration of Olinone had no such effect (Figure 2B). JQ1 reverted TGFβ1-stimulated αSMA expression to the unstimulated basal level. A lesser RVX208 effect than JQ1 on αSMA expression was likely due to their different BD2-binding affinities (~500 nM vs 50 nM, respectively)^15^.

**Figure 2.**
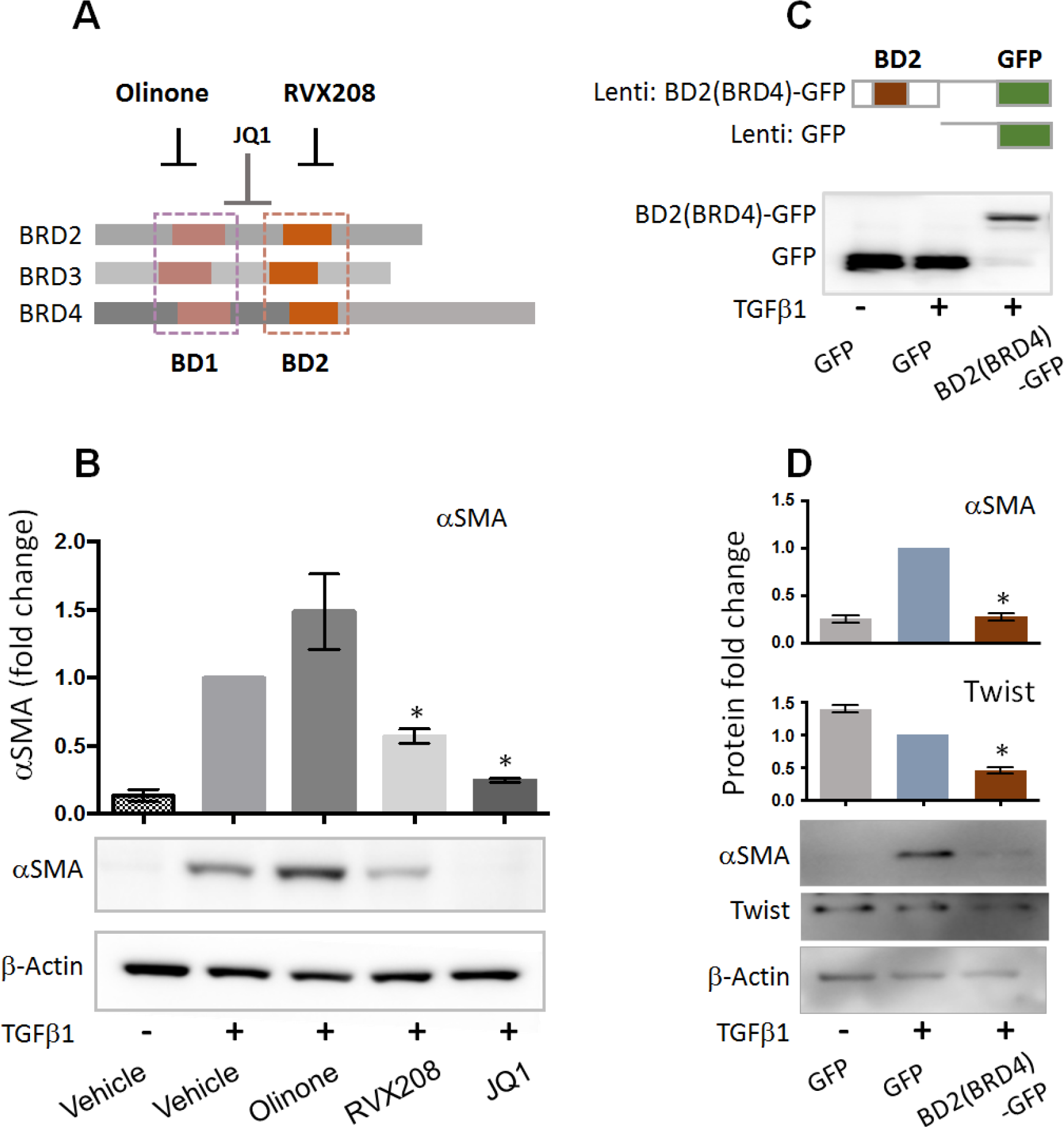
Disruption of BET bromodomain 2 (BD2) but not that of BD1 mitigates TGFβ1-induced EndoMT. A. Schematic: Selectivity of BET bromodomain blockers; Olinone for BD1, RVX208 for BD2, and JQ1 for both, in all BETs. B. Effects of BD-selective BET inhibitors on αSMA as a canonical EndoMT marker. Rat aortic ECs were cultured, starved, pre-treated with vehicle or a BET inhibitor (5 μM Olinone, 5 μM RVX208, or 1 μM JQ1), and then stimulated with TGFβ1, as described in Figure 1A. After 96h TGFβ1 stimulation, the cells were harvested for Western blotting assay of αSMA. C. Overexpression of dominant-negative BRD4-specific BD2 domain. BD2(BRD4) fused with GFP was expressed in ECs using a lentivirus and detected with a GFP antibody. D. Expression of BD2(BRD4) represses TGFβ1-induced EndoMT. ECs were transduced for 6h with a lentivirus of GFP control or BD2(BRD4)-GFP in the complete medium, recovered in fresh same medium for 24h, and then starved for 6h in the basal medium containing 1% FBS before TGFβ1 stimulation (96h) and Western blotting. Quantification of densitometry reading (normalized to β-actin): For each independent repeat experiment, the control value (vehicle or GFP+TGFβ1) is presented as 1, and the values from other conditions are normalized as fold changes. Normalized values from ≥ 3 repeats are averaged (mean ± SEM); *P<0.05 compared to the normalizing control.

Given the predominant role of BRD4 in EndoMT (Figure 1C) and the evidence for BD2 accounting for the bromodomain effect on EndoMT in all BETs (*vs* BD1, Figure 2B), we investigated if the BD2 domain in BRD4 mainly accounted for BRD4’s action in directing EndoMT. To address this, we utilized a lentiviral vector to express the BRD4-specific BD2 domain (Figure 2C) that serves as a dominant-negative competitor to endogenous BD2, as previously established ^16^. Indeed, compared to the empty vector (GFP control), expression of GFP-tagged BD2(BRD4) completely abolished TGFβ1 stimulation of αSMA protein expression (Figure 2D) while also significantly reducing Twist expression. Together, these results support BD2 as determining the action of BRD4 in TGFβ1-induced EndoMT and suggest targeting BD2 in BRD4 would have the greatest impact on EndoMT.

### BRD4 silencing averts TGFβ1-stimulated ZEB1 protein up-regulation

ZEB1 has been recently identified as a key transcription factor governing transcription programs for both EMT and EndoMT^1, 17^. In addition, Snail and Twist have also been shown to be important for EMT, although their differential and/or cooperative functional relationships with ZEB1 remain to be understood ^13^. Given BETs as an epigenetic mechanism for coupling transcription factors to gene expression programs and the data presented for BET regulation of EndoMT, we next investigated BETs involvement in controlling these novel transcriptional mediators of EndoMT. TGFβ1 increased ZEB1 protein, which was completely repressed in the presence of JQ1, reducing this protein to non-stimulated basal levels (Figure 3A). We then used siRNAs to dissect the roles of specific BET family members. While BRD4 siRNA abolished TGFβ1-stimulated increases in ZEB1 protein, siRNA to either BRD2 or BRD3 had no significant effect on ZEB1 (Figure 3B). Importantly, all three siRNAs had similar efficiency in repressing their respective BET member (Figure 1B), thus eliminating variable knockdown as an explanation for these effects. Although TGFβ1 did not increase Snail and Twist protein levels, both JQ1 (Figure 3A) and BRD4 siRNA (Figure 3B) markedly reduced Snail and Twist protein levels whereas BRD2 and BRD3 siRNAs had no significant effects. These results indicate BETs, and more specifically, BRD4, as a mechanism controlling the expression of ZEB1, a central mediator in EndoMT.

**Figure 3.**
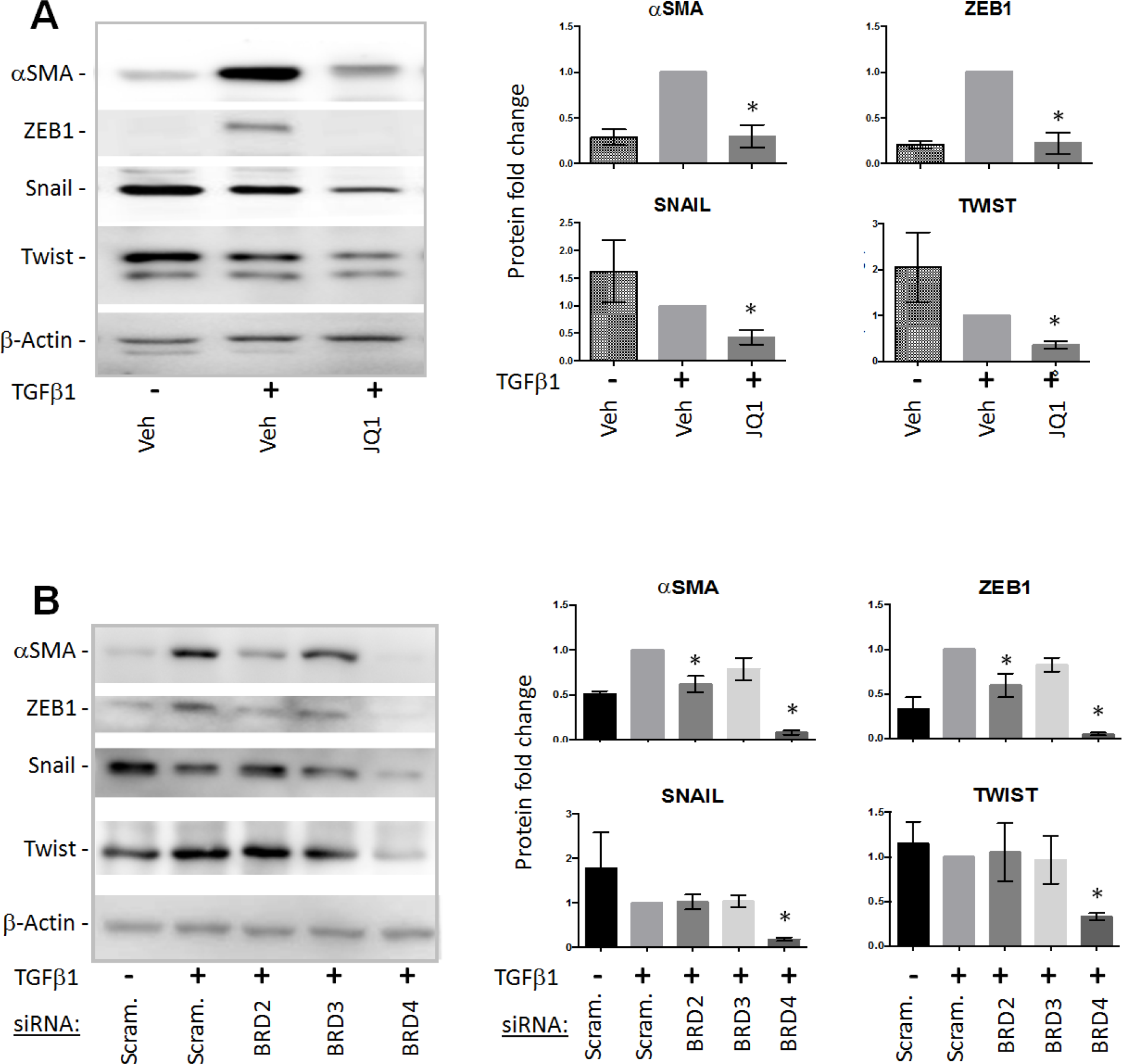
BET inhibition blocks TGFβ1-induced increase of the EndoMT-associated master transcription factor ZEB1. **A.** Effect of pan-inhibition of BETs with JQ1. Experiments (JQ1 pre-treatment and TGFβ1 stimulation) were performed as described in Figure 1A. At 96h after TGFβ1 stimulation, ECs were harvested for Western blotting analysis of αSMA and EndoMT-associated transcription factors. **B.** Effect of knockdown with an siRNA specific for BRD2, BRD3, or BRD4. Experiments (siRNA transfection and TGFβ1 stimulation) were performed as described in Figure 1C except that cells were harvested for Western blotting at 96h after TGFβ1 stimulation. Quantification of Western blots was performed as described for Figure 2; mean ± SEM; n ≥ 3; *P<0.05 compared to the normalizing control.

### Lentiviral transgene of BRD4 shRNA or dominant-negative BRD4-specific BD2 mitigates neointima formation in rat vein grafts

Given recent evidence for EndoMT as being critical to neointima formation ^1, 3^, our data for BETs, and more specifically BRD4 regulation of mediators and markers of EndoMT suggested BRD4 manipulation would decrease neointima formation in vein grafts *in vivo*. To pursue this, we employed an established cuff technique ^18^ in a rat vein graft model to investigate specific BRD2, BRD3 and BRD4 involvement in neointimal thickening (Figure 4). This vein graft model allowed for uptake of BRD-specific shRNA-expressing (or control) lentivirus by the intact endothelium of rat jugular vein explants (Figure S2). The explants were incubated with lentivirus for 4 hours prior to their grafting onto carotid arteries (diagrammed in Figure 4A). Vein grafts were harvested and analyzed 4 weeks later, including measurement of intima/media area ratio (I/M) as a measure of neointimal lesion formation ^7^. Lentiviral transduction of either BRD2 or BRD3 shRNA in vein graft explants had no effect on I/M ratio as compared to empty lentiviral control vectors (Figure S3). However, in striking contrast, lentiviral BRD4 shRNA transduction significantly reduced the I/M ratio by ~65% as compared to control vectors (Figure 4, B and C). Furthermore, lentiviral expression of dominant-negative BD2(BRD4) also effectively mitigated neointima formation, as indicated by a 78% decrease of the I/M ratio compared to control (Figure 4, D and E). These results provide *in vivo* evidence that BRD4, but not BRD2 or BRD3, and more specifically, the BD2 domain in BRD4, critically mediates neointima formation in the rat vein graft model, findings consistent with our *in vitro* data (Figures 1-3).

**Figure 4.**
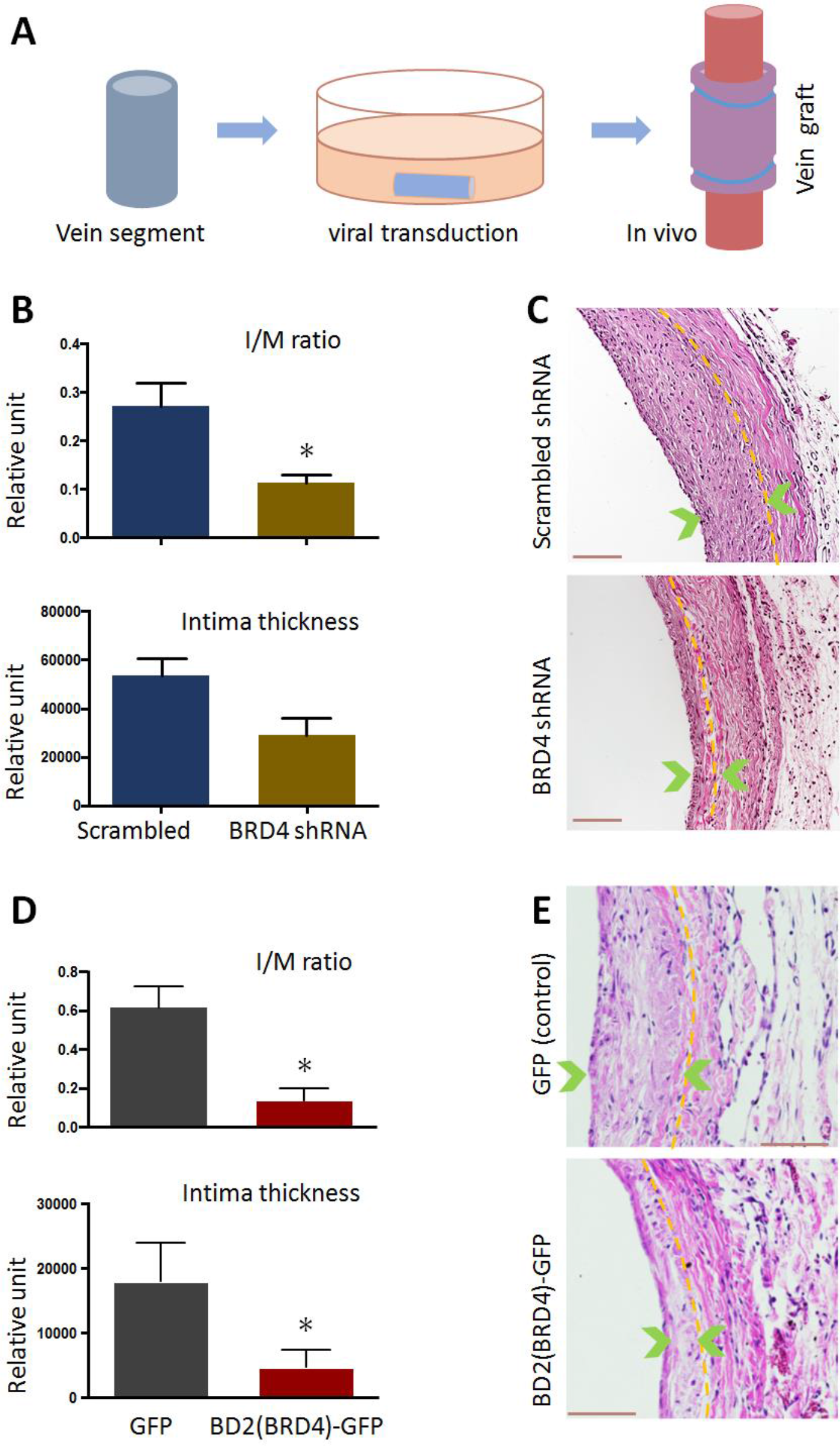
Transduction of BRD4 shRNA or dominant-negative BD2(BRD4) into vein grafts inhibits neointimal formation *in vivo*. *Ex vivo* lentiviral transduction of rat jugular vein segments and their grafting to carotid arteries are described in detail in Methods. Vein grafts were harvested 4 weeks after grafting; cross-sections were H&E-stained for analysis of the intima/media (I/M) area ratio. **A.** Diagram of *ex vivo* lentiviral transduction of a rat jugular vein followed by grafting to a carotid artery. A cuff-assisted suturing (non-stitching) technique was applied. **B** and **D.** Quantified morphometric data indicating reduced I/M ratio in the vein grafts transduced with a lentiviral vector to express BRD4 shRNA or dominant-negative BD2(BRD4), respectively. Mean ± SEM; n = 6 rats; *P<0.05. **C** and **E.** Representative H&E-stained cross-sections of vein grafts (in B and D, respectively). The thickness of neointima is indicated between green arrowheads; dashed line marks the internal elastic lamina (IEL). Scale bar: 100 μm.

## Discussion

We provide *in vitro* and *in vivo* evidence here for the BET bromodomain protein BRD4 as the specific BET family member that controls the EndoMT process, doing so by coordinating the endothelial transcriptional program induced by TGFβ1, including established EndoMT markers like αSMA as well as novel transcription factors like ZEB1. *In vitro*, BRD4 (*vs* BRD2 or BRD3) silencing is predominant and sufficient to block EndoMT, with this BRD4 effect dependent on BD2 but not BD1. *In vivo*, either BRD4 knockdown or dominant-negative BD2(BRD4) expression represses neointima formation, a key step in vein graft failure involving EndoMT.

Although an intervention commonly employed to relieve ischemia, vein grafting is a flawed strategy, with almost half of grafts failing in 4 years or less due primarily to flow-limiting neointima formation^3^. As a result, intense interest has focused on neointima formation and thus far unsuccessful attempts to modulate this vein graft response. Recent studies highlight EndoMT as a novel, fundamental driver of neointima formation in vein grafts as well as atherosclerosis ^2, 3^. During EndoMT, the endothelium undergoes transcriptional reprogramming and cell state transition, losing endothelial properties and expressing SMC-like markers. Although the identification of EndoMT has been a significant advance, thus far, no clear targets for therapeutically modifying EndoMT responses have emerged.

BETs have recently emerged as epigenetic regulators that control differentiation and cell state transitions, doing so by localizing master transcription factors and core transcriptional machinery at multiple discrete, marked chromatin sites through BET binding to acetylated lysines on histone tails. For various reasons, BET targeting is being pursued intently in oncology, including BET rearrangements as a direct cause of some cancers, BET involvement in defining specific gene cassettes underlying myeloma, and the development of specific BET inhibitors with favorable pharmacokinectic profiles ^5, 12^. More recent studies from us and others have strongly implicated BETs in cardiovascular responses involving cell state transitions, with BET inhibition blocking cardiac hypertrophy in response to trans-aortic constriction ^8^, the TNFα-induced NFκB program in ECs and atherosclerosis ^4^, and SMC proliferation/migration ^7^, among others. Given these findings, we hypothesized BETs as potential novel determinants of the unique cell state transition/trans-differentiation found in EndoMT and subsequent neoinitima formation, as borne out here. While the common use of pan-BET inhibitors like JQ1 generally precludes elucidation of relative BRD2, BRD3 or BRD4 contributions to vascular responses, we establish here that EndoMT depends on BRD4, and more specifically, the BD2 of BRD4.

Consistent with a prominent effect of BET inhibition on blocking EndoMT observed herein^11^, JQ1 has been reported to mitigate EMT in carcinomic ^17, 19^ and pulmonary^20^ epithelial cells, although without identification of the specific BET family members or BDs involved. Interestingly, a very recent report identified BRD2 as the key regulator of EMT with BRD3 and BRD4 playing an opposite role ^21^. This study, which contrasts with our finding of EndoMT being dependent on BRD4, may align with other studies identifying BRD4 action as varying as a function of cell-specific as well as stimulus-dependent contexts ^6^. Examples include non overlapping inflammatory gene regulation in macrophages ^22^ versus ECs ^4^, Myc regulation in myeloma ^23^ but not cardiomyocytes ^8^ and in stimulated T cells ^24^. Alternatively, experimental issues may underlie observed differences in BET responses. While TGFβ1 was used to stimulate EndoMT here, no specific stimulus was used in investigating BET involvement in breast cancer EMT ^21^. Another factor in dominant BRD4 versus BRD2 action may involve our studies being done in primary rat aortic ECs not immortalized cancer cell lines. Finally, EndoMT and EMT are overlapping yet distinct processes, with likely both shared as well as distinct regulatory mechanisms.

The potent role of BETs in controlling transcriptional programs in differentiation as well as pathologic states like cancer and atherosclerosis has prompted interest in better understanding BET action. Our data provides new insight into fundamental aspects of BET biology in vascular cells, with evidence that BET regulation of EndoMT depends on the second (BD2) but not the first (BD1) of the two tandem bromodomains present in BRD2, BRD3 and BRD4 ^6^. BRD4 is unique in possessing a C-terminal domain, which has a reported critical role in co-activating transcription ^5^. While BRD4 is recruited to specific gene loci via bromodomain binding to acetylated sites, the interaction of its C-terminal domain with the transcription elongation factor (pTEFb) facilitates RNA polymerase II activation. Whether the C-terminal domain present in BRD4 but absent in BRD2 ^5^, explains BRD4’s more dominant role (*vs* BRD2) in directing the EndoMT program requires further study. Similarly, whether BD1 might be more important than BD2 in other cell types or settings will require additional investigation including use of approaches employed here. Despite their significant sequence homology and structural similarity^12^, our data highlights distinct effects mediated through BD2 versus BD1, variations that may have functional importance given the prospect of selective BD targeting for therapeutic purposes, as has been raised. ^14, 15 5^.

The EndoMT cell state transition can be viewed as a form of cellular reprogramming, with parallels to physiologic processes like differentiation and also pathologic issues like oncogenic transformation. Our identification of BRD4 as coordinating a TGFβ1-induced EndoMT transcriptional program provides new insight into the burgeoning interest in this mesenchymal transition. We show BRD4 regulation orchestrates changes in ZEB1, Snail, and Twist, downstream mediators of EndoMT ^1^. Although not previously studied in terms of vascular biology, BRD4 has been shown to interact with Twist on chromatin in breast cancer cells ^13^. We show TGFβ1-induced EndoMT increased ZEB1 protein but reduced Snail and Twist levels, with either the BET inhibitor JQ1 or BRD4 silencing abolishing these responses and preventing changes in EndoMT phenotypic markers like αSMA (Figures 1 and 3). Of note, it has been recently identified that metabolic factors, such as endothelial fatty acid oxidation (FAO), regulate TGFβ1-induced EndoMT as well ^25^. Nevertheless, our findings argue for further consideration of BET involvement in transcriptional regulation of EndoMT.

## Conclusions

By dissecting the functional roles of BET family members and BD domains, we reveal here that BRD4, and more specifically, the BRD4-specific BD2, plays the predominant role in TGFβ1-induced EndoMT, a key process in vein graft failure. As seen in other settings, we show BRD4 coordinates changes of transcription factors upon cell state transitions. These findings reveal novel mechanisms involved in EndoMT and BRD4 action, underscore selective BET family member action, and have implications for ongoing efforts at BET inhibition for therapeutic purposes, including support for selective BET bromodomain effects. Our results also suggest inhibition of BD2 in BRD4 as a possible strategy for the unmet need of preventing vein graft failure.

## Acknowledgement

This work was supported by NIH grants R01 HL133665 (to L.-W. G.) and R01HL-143469 (to K.C.K., L.-W. G. and S.G.), and an AHA pre-doctoral award 17PRE33670865 (to M.X.Z.).

