## Supplemental figures and tables for "The BD2 domain of BRD4 is a determinant in EndoMT and vein graft neointima formation"

*Short title: Role of BRD4 in EndoMT and vein graft neointima*

* Corresponding author:

Lian-Wang Guo, Ph.D.

Department of Surgery and Department of Physiology & Cell Biology
The Ohio State University, Columbus, OH 43210, USA

#### Methods

##### ***Rat aortic EC culture, induction of EndoMT, and pretreatment with BET inhibitors***

Rat primary aortic endothelial cells (ECs) were purchased from Cell Biologics (Cat.RA-6052). For cell expansion, the culture was maintained at 37°C/5% $CO_2$ in the Complete Rat Endothelial Cell Medium containing growth factor supplement and 2% FBS (Cell Biologics, Cat. M1266); Accutase (Lifetechnologies, Carlsbad, CA) was used for cell detachment. The cells used for experiments were at passages 5-7. For induction of EndoMT, cells were starved in the Endothelial Cell Culture Basal Medium (Cell Biologics, Cat. M1266b) with 1%FBS for 6h, and TGFβ1 (human recombinant, R&D Systems Inc., MN) was added to a final concentration of 50 ng/ml and incubated for 72h or 96h before cell harvest for mRNA or protein determination respectively. In the experiments using BET inhibitors, cells were pretreated with JQ1, RVX208 (Apexbio, Houston, TX, USA), or Olinone (Cat. GLXC-05021, Glixx Laboratories, Southborough, MA, USA) at indicated concentrations, or vehicle control (equal volume of DMSO, Sigma-Aldrich, St. Louis, MO) for 2h prior to induction of EndoMT with TGFβ1.

***Quantitative real-time PCR (qRT-PCR) to determine mRNA levels***

The assay method was described in our previous report [1]. Briefly, total RNA was isolated from cultured cells using the Trizol reagent (Invitrogen, Carlsbad, CA). Potential contaminating genomic DNA was removed by using gDNA Eliminator columns provided in the kit. RNA was quantified with a Nanodrop NP-1000 spectrometer, and RNA of 1 µg was used for the first-strand cDNA synthesis (Applied Biosystems, Carlsbad, CA). Quantitative RT-PCR was then performed using Quant Studio 3 (Applied Biosystems, Carlsbad, CA). The house keeping gene GAPDH was used for normalization. Each cDNA template was amplified in triplicate PerfeCTa SYBR® Green SuperMix (Quantabio) with gene specific primers, as listed in Table S1.

***In vitro knockdown of BET family members with siRNAs***

ECs were grown to 70% confluency in 6-well plates in the Complete Rat Endothelial Cell Medium with growth factor supplement. The BRD2-, BRD3-, or BRD4-specific siRNA was added to transfect ECs for overnight using the RNAi Max reagent (Thermo Fisher, cat.13778150). Cells were then recovered in the complete medium for 24h. If EndoMT was induced, the culture was changed to the basal medium containing 1% FBS and incubated for 6h prior to TGFβ1 stimulation. The siRNAs were ordered from Thermo Fisher (sequences listed in Table S2).

***Adeno- and lenti-viral vectors for in vitro and ex vivo transduction***

Adenovirus stocks for HA-tagged BRD4 (AdBRD4-HA, cat# 073066A) and empty vector control (AdNull, cat# 000047A) were purchased from Applied Biological Materials (Canada). An MOI of 100 was used for in vitro infection of primary rat aortic ECs. Cells were grown to 70% confluency in 6-well plates in the Complete Rat Endothelial Cell Medium with growth factor supplement, and adenovirus was added and incubated for 6h followed by 24h recovery in the complete medium. Cells post infection were incubated in the basal medium (1% FBS) for 6h and then treated with TGFβ1 for 96h before harvest for Western blotting analysis.

The piLenti-shRNA-GFP vectors for the production of a scrambled shRNA or an shRNA specific for rat BRD2, BRD3, or BRD4 were purchased from Applied Biological Materials (Canada). The most efficient sequences (of siRNAs, final products) are as follows: BRD2, CCACAATGGC TTCTGTACCA; BRD3, AGTGAGTGTAT GCAGGACTTCAACACCAT; BRD4, GCGAATCTAGCTCCTCTGACAGTGAAGAC. The lentiviruses were packaged in Lenti-X 293 cells (Clontech,cat#632180) using a three-plasmid expression system (piLenti-siRNA-GFP, psPAX2 and pMD2.G) as we recently reported[1].

The lentiviral construct for the expression of GFP fusion with BRD2 or BRD4 or BD2(BRD4) was generated using a GFP-expressing lenti-vector (empty vector, control) that was kindly provided by Dr. Ming-Liang Chu (Guizhou Renmin Hospital, China). Molecular cloning and lentivirus packaging in Lenti-X 293 cells were performed following our previously reported method [1]. Briefly, before harvesting viruses, a minimum of 5 x 105 infection unit (IFU)/ml in supernatant was guaranteed using Lenti-x GoStix (Takara, cat.631243). The crude viral solution was then concentrated using Lenti-x concentrator (Takara, cat.631232) to a final concentration of 108-109 IFU/ml using the Lenti-x qRT-PCR Titration Kit (Takara, cat.631235). An MOI of 10 was used for in vitro infection of primary rat aortic ECs with lentiviruses to express GFP alone (empty vector), BRD2-GFP, BRD4-GFP, or BD2(BRD4)-GFP. Cells were grown to 70% confluency in 6-well plates in the Complete Rat Endothelial Cell Medium with growth factor supplement (Cell Biologics, Cat. M1266), and lentivirus plus polybrene (Santa Cruz.,Cat.sc-134220) was added and incubated for 6h followed by 24h recovery in the complete medium. Cells post infection were incubated in the basal medium (1% FBS) for 6h and then treated with TGFβ1 for 96h before harvest for Western blotting analysis. The use of lentiviruses for in vivo experiments is described below.

##### *Western blotting to determine changes of protein levels*

Cells were collected and lysed in RIPA buffer (Thermo Fisher, cat.89900) on ice, and a protease inhibitor cocktail (Thermo Fisher, cat.87785) was included. Cell lysates were quantified for protein concentrations using the Bio-Rad DC™ Protein Assay kit (cat.5000112) and loaded to a 10% SDS-PAGE gel. The transfer to a PVDF membrane, incubations with primary and secondary antibodies were performed following our established methods [1]. The antibody sources and dilution ratios are listed in Table S3. Specific protein bands on the blots were illuminated by applying enhanced chemiluminescence reagents (Pierce), recorded by an Azure C600 Imager (Azure Biosystems), and quantified using NIH Image J.

##### *Ex vivo gene transfer and rat jugular vein graft model*

All animal studies conform to the *Guide for the Care and Use of Laboratory Animals* (National Institutes of Health) and protocols approved by the Institutional Animal Care and Use Committee at The Ohio State University. Male Sprague-Dawley rats (Charles River; 300–330 g) were used.

The model of rat autologous external jugular vein to common carotid artery reversed graft interposition was performed with modifications based a method previously described for mice[2]. Briefly, prior to surgery, rats were anesthetized with isoflurane (5% for inducing and 2.5% for maintaining anesthesia). After a midline incision, the posterior facial branch of the external jugular vein was exposed by aseptic incision in the ventral neck. An 8 mm segment of the vein was harvested and temporarily placed in cold Hank's Balanced Salt Solution (HBSS, containing $Ca^{2+}$ and $Mg^{2+}$) until use for ex vivo gene transfer.

For knockdown of BRD2, BRD3, or BRD4, or for overexpression of BD2(BRD4)-GFP in vein grafts, ex vivo lentiviral transduction was performed[3]. Briefly, prior to grafting, jugular vein segments (explants) were incubated for 4 h (37°C/5% $CO_2$) with the lentivirus expressing BRD2 (or 3 or 4)-specific shRNA or BD2(BRD4)-GFP in the RPMI 1640 medium containing viral particles (>107 IFU/ml) and heparin (10 U/ml).

For vein grafting, the right common carotid artery was dissected and severed between two ligatures generating two ends. Each end of the carotid artery was clamped together with an overwrapped plastic cuff and the ligature was removed. Each end of the carotid artery was then everted over the cuff and secured with a suture, and a vein explant (after viral transduction) was sleeved over both everted arteries on cuffs and secured with sutures.

To prevent thrombosis, prior to vein graft interposition, heparin s.c. was injected (300 U/kg body weight), and the lumens of carotid arteries and vein grafts were periodically flushed with heparinized saline during the grafting procedure. Clamps were then removed to restore blood flow, and skin wounds were closed using clips. Following subcutaneous injection of heparin (200 U/kg), animals were recovered and returned to husbandry. Rats were euthanized at 4 weeks post vein grafting, and the grafts were collected for histology analysis.

##### *Morphometric analysis of intimal hyperplasia (IH)*

At 4 weeks after the grafting surgery, the whole vein graft except the perianastomotic regions was harvested and immersed in 4% paraformaldehyde for 24 h and then paraffin-embedded. Paraffin sections (5 μm thick) were excised from the grafts at equally spaced intervals and then H&E-stained for morphometric analysis. Planimetric parameters as follows were measured on the sections and calculated using Image J as described in our previous report [1]: area inside external elastic lamina (EEL area), area inside internal elastic lamina (IEL area), lumen area, intima area (= IEL area- lumen area), and media area (= EEL area – IEL area). Intimal hyperplasia was quantified as a ratio of intima area versus media area (I/M). Measurements were performed by a student blinded to the experimental conditions. There were two animal groups each including 5-6 rats: one group grafted with a vein that was transduced with lentivirus to express scrambled shRNA (or GFP) control, and the other group with a vein transduced with lentivirus to express BRD4 shRNA (or BRD4-specific BD2). The data from all 4-6 sections were pooled to generate the mean for each animal. The means from all the animals in each treatment group were then averaged, and the standard error of the mean (SEM) was calculated.

##### *Statistical analysis*

To confirm one result, at least three independent repeat experiments were performed. Data are presented as mean ± SEM. Two-condition comparison was analyzed with Student's t test using Prism version 4.0 (GraphPad Software). Multi-condition comparison was analyzed with one-way ANOVA followed by Bonferroni post hoc test. Significance was set to $P < 0.05$.

#### Supplemental figures

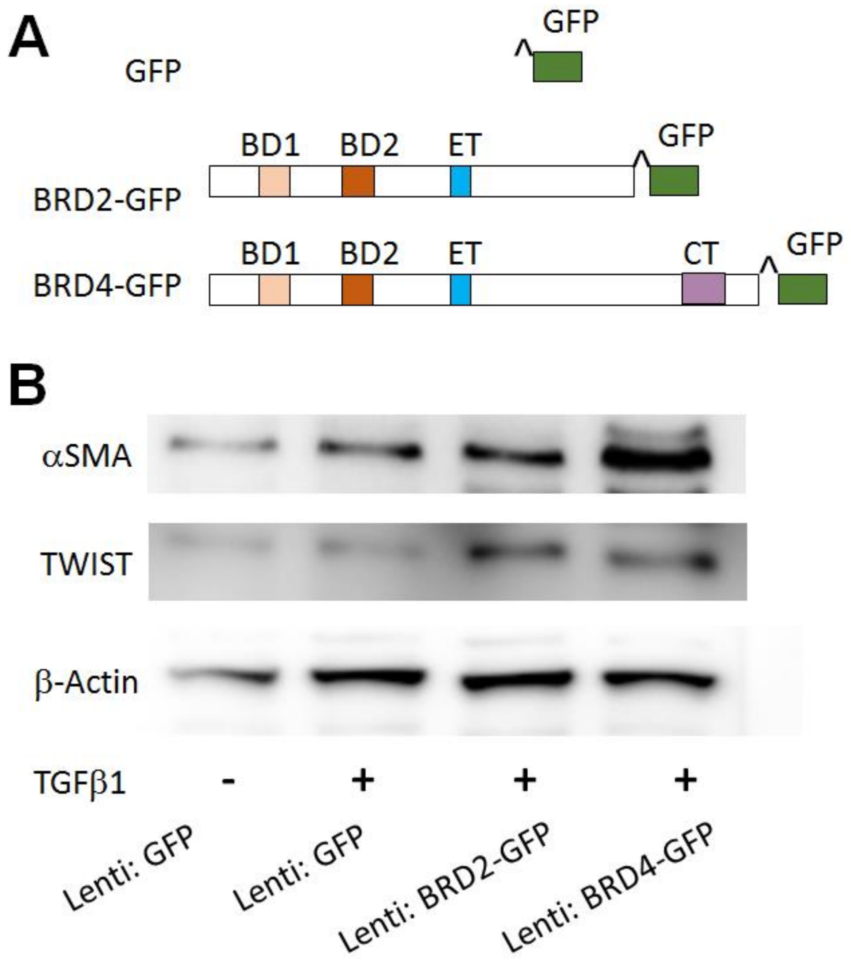

**Figure S1. Gain-of-function expression of BRD2 and BRD4**

Experimental conditions were the same as described for Figure 1D except that lentiviruses were used for Figure S1. Briefly, ECs were transduced with empty vector (GFP control) or GFP-fused with BRD2 or BRD4 (diagramed in A). ET, extraterminal domain; CT, C-terminal domain. Shown in B are representative Western blots of two similar experiments.

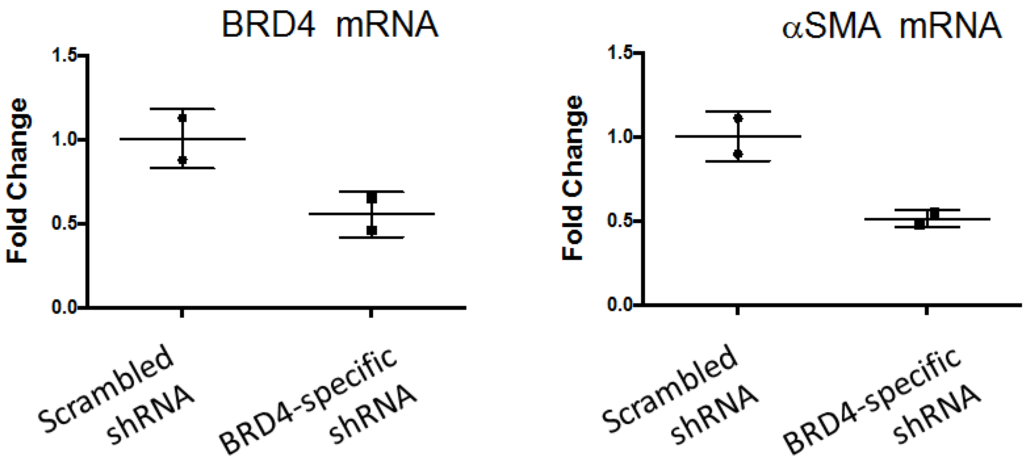

**Figure S2. In vivo BRD4 knockdown efficiency**

Ex vivo transduction of vein explants with lentivirus expressing BRD4 shRNA or scrambled shRNA (control) and vein grafting were performed as described in Figure 4. The vein grafts were collected 8 days after the grafting surgery and the homogenates were used for qRT-PCR determination of BRD4 or αSMA mRNA levels. Data were obtained from two separate experiments using two rats.

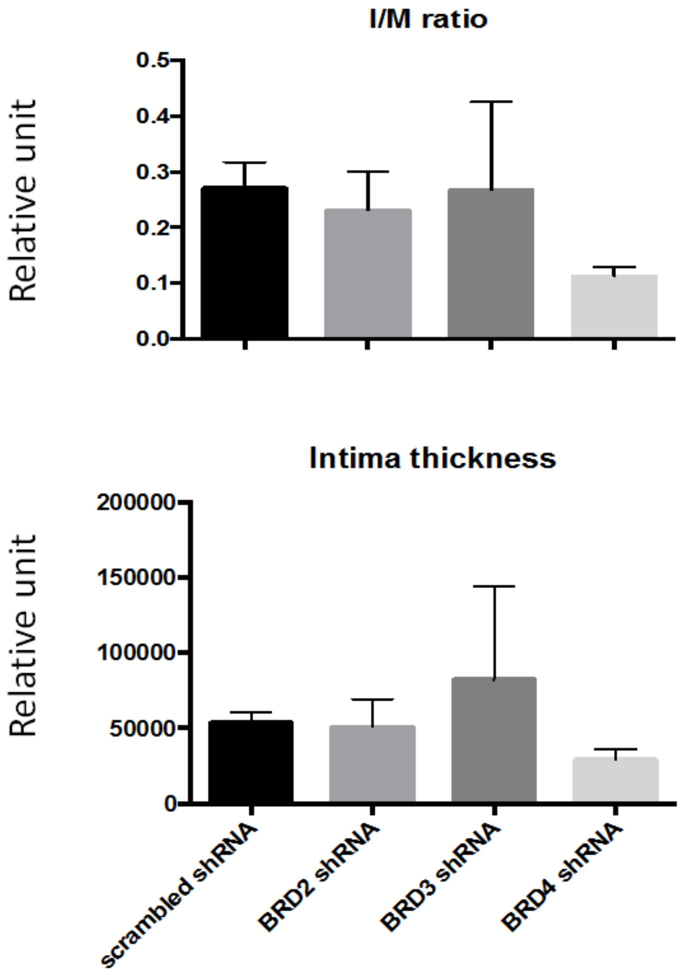

**Figure S3. Effect of shRNAs of BRD2, BRD3, and BRD4 on vein graft neointima**

Ex vivo transduction of vein explants with lentivirus expressing BRD2 shRNA or BRD3 shRNA or scrambled shRNA, vein grafting, and morphometric analysis were all performed in parallel to the experiments with BRD4 shRNA-expressing lentivirus. Experimental conditions were the same as described for Figure 4.

### Supplemental Tables

**Table S1. Primers used for qRT-PCR**

| | Forward | Reverse |
|---|---|---|
| $\alpha$SMA (ACTA2 | ACCTTCAATGTCCCTGCCATGTA | ACGAAGGAATAGCCACGCTCA |
| Calponin (CNN1) | CGGCGTCACCTCTATGATCC | AGCGTGTCACAGTGTTCCAT |
| Collagen-I (COL1A1) | ATCAGCCCAAACCCCAAGGAGA | CGCAGGAAGGTCAGCTGGATAG |
| BRD2 | GGGTCTGCCGGATTATCACA | GCCCCCTTCTTATGGCTGTT |
| BRD3 | AAGATGGTGAGGTCCCACAG | GGTACTCACGGCTGTCCATT |
| BRD4 | CTGCCAGTAATGGGGGATGG | ATTGGTGCTGGCTGCATTTG |
| GAPDH | CCCCATTTGATGTTAGCGGG | CCCCATTTGATGTTAGCGGG |

**Table S2. siRNAs for knocking down BET family members**

| | sense | Anti-sense |
|---|---|---|
| BRD2 | GCUUGAACGAUACGUUUUA | UAAAACGUAUCGUUCAAGC |
| BRD3 | AGGAAACCAUUGUCAACAATT | UUGUUGACAAUGGUUUCCUCT |
| BRD4 | GCAUCAACUUCUCCGCAGATT | UCUGCGGAGAAGUUGAUGCTT |

**Table S3. Antibodies for Western blotting**

| Antibody | Company | Cat# | Dilution |
|---|---|---|---|
| Anti-rabbit $\alpha$SMA monoclonal | Abcam | ab32575 | 1:1000 |
| Anti-mouse ZEB1 monoclonal | Santa Cruz | sc-515797 | 1:200 |
| Anti-rabbit BRD4 monoclonal | Abcam | Ab128874 | 1:1000 |
| Anti-mouse SNAIL monoclonal | Santa Cruz | sc-271977 | 1:200 |
| Anti-mouse TWIST monoclonal | Novus Biologicals | 10E4E6 | 1:1000 |
| Anti-mouse $\beta$-actin monoclonal | Abcam | Ab8226 | 1:1000 |
| Anti-rabbit GFP polyclonal | CST | 2956 | 1:1000 |
